# Bioinformatic and fine-scale chromosomal mapping uncover the essence and evolution of eliminated chromosomes in the Japanese hagfish, *Eptatretus burgeri*, through repetitive DNA family analysis

**DOI:** 10.1101/2023.05.28.542657

**Authors:** Kohei Nagao, Yoshiki Tanaka, Rei Kajitani, Atsushi Toyoda, Takehiko Itoh, Souichirou Kubota, Yuji Goto

## Abstract

In the Japanese hagfish, *Eptatretus burgeri*, approximately 21% of the genomic DNA in germ cells (2n=52) consists of 16 chromosomes (eliminated [E]-chromosomes) that are eliminated from presumptive somatic cells (2n=36). To uncover the eliminated genome (E-genome), we have identified 16 eliminated repetitive DNA families from eight hagfish species, with 11 of these repeats being selectively amplified in the germline genome of *E. burgeri*. Furthermore, we have demonstrated that six of these sequences, namely EEEb1–6, are exclusively localized on all 16 E-chromosomes. This has led to the hypothesis that the eight pairs of E-chromosomes are derived from one pair of ancestral chromosomes via multiple duplication events over a prolonged evolutionary period. NGS analysis has recently facilitated the re-assembly of two distinct draft genomes of *E. burgeri*, derived from the testis and liver. This advancement allows for the prediction of not only nonrepetitive eliminated sequences but also over 100 repetitive and eliminated sequences, accomplished through K-mer-based analysis. In this study, we report four novel eliminated repetitive DNA sequences (designated as EEEb7–10) and confirm the relative chromosomal localization of all eliminated repeats (EEEb1–10) by fluorescence *in situ* hybridization (FISH). With the exception of EEEb10, all sequences were exclusively detected on EEEb1-positive chromosomes. Surprisingly, EEEb10 was detected as an intense signal on EEEb1-positive chromosomes and as a scattered signal on other chromosomes in germ cells. The study further divided the eight pairs of E-chromosomes into six groups based on the signal distribution of each DNA family, and fiber-FISH experiments showed that the EEEb2–10 family was dispersed in the EEEb1-positive extended chromatin fiber. These findings provide new insights into the mechanisms underlying chromosome elimination and the evolution of E-chromosomes, supporting our previous hypothesis.

## Introduction

In multicellular organisms, the genomic information in all cells is generally identical throughout their development and differentiation. However, certain species display differences between somatic and germline genomes as a result of genomic rearrangements. This phenomenon, called programmed genome rearrangement (PGR), is responsible for the expulsion of genomic regions that are identical in all somatic cells. PGR was first reported by Boveri in the nematode *Parascaris equorum* [1]. In this species, the fertilized egg and presumptive germ cells retain a pair of large chromosomes (2n=2). However, during the second to sixth cleavages, PGR leads to the fragmentation of these chromosomes in blastomeres, followed by their elimination through nondisjunction at the M phase. Consequently, the fragmented chromosomes differentiate into somatic cells. Chromatin diminution/chromosome elimination refers to PGR, where either chromosome fragments or chromosomes are eliminated during early embryogenesis; PGR has been reported in nematodes, ciliates, insects, crustaceans, and vertebrates [2, 3].

PGR in vertebrates was first discovered in the Japanese hagfish, *Eptatretus burgeri*. In this species, 52 chromosomes including 36 C-band negative and 16 C-band positive chromosomes were observed in the spermatogonial cells, whereas only 36 C-band negative chromosomes were retained in the somatic cells, indicating the exclusive elimination of C-band-positive chromosomes [4]. In that study, quantification of the nucleic DNA in both cells clearly revealed that approximately 20.9% of the germline genomic DNA is eliminated in the somatic cells. Further intensive studies have confirmed chromosome elimination in seven hagfish species (*E. stoutii*, *E. okinoseanus*, *E. cirrhatus*, *Paramyxine atami*, *P. sheni*, *Myxine glutinosa,* and *M. garmani*). In these species, 2 to 62 chromosomes are shed from presumptive somatic cells, corresponding to 20.9–74.5% of the germline genomic DNA [5]. Moreover, molecular genetic analyses revealed that 16 highly repetitive DNA families are selectively amplified and eliminated sequences [6–11]. Four eliminated repetitive DNA families, namely EEEo1 (Eliminated Element of *E. okinoseanus* 1), EEEo2, EEPa1, and EEEb2, are conserved in the germline genomes of the hagfish species studied [6–10, 12]. Of the 16 eliminated DNA families in *E. burgeri*, 11 (EEEb1, EEEb2, EEEb3, EEEb4, EEEb5, EEEb6, EEEo1, EEEo2, EEPa1, EEPs1, and EEPs4) were selectively amplified in the germline genome, accounting for about 43.3% of the eliminated DNA, as determined by Southern- and slot-blot assays. A fluorescence *in situ* hybridization (FISH) assay also showed that six eliminated DNA families, EEEb1–6, are exclusively localized on all eliminated chromosomes (E-chromosomes) with quite similar signal distributions [11]. Hence, we hypothesized that the eight pairs of E-chromosomes are derived from a single pair of ancestral chromosomes that underwent multiple duplication events caused by a meiotic drive over a long evolutionary period.

In the past decade, whole-genome sequencing (WGS) has been carried out in several animals undergoing PGR [13–22]. Comparative genome analysis between the germline and somatic genomes has revealed that not only numerous repetitive sequences but also hundreds of protein-coding genes, some of which appear to be involved in the developmental process, are eliminated from the presumptive somatic cells. One of the most extensively studied animals in the context of chromatin diminution is the ascarid nematode, *Ascaris suum.* The complete genome assemblies of the germline and somatic genome have been constructed, demonstrating that 24 germline chromosomes (19 autosomes and five sex chromosomes) break into 36 somatic chromosomes (27 autosomes and nine sex chromosomes) through 72 chromosomal breaks, with the excision of 12 interstitial regions and the loss of 48 telomere-subtelomere regions [15, 23, 24]. The mapping of homologous gene sequences between *A. suum* and *Caenorhabditis elegans* demonstrated that the chromosomal organization of these species is largely conserved, although it should be noted that this conservation is accompanied by various instances of chromosomal fissions and fusions. Thus, WGS can be a powerful tool for the complete elucidation of released sequences and the consideration of their evolutionary processes.

Recently, the complete eliminated genome (E-genome), consisting of nonrepetitive sequences and genes, was elucidated through comparative genome analysis. This was accomplished by assembling WGS derived from germ cells (G-genome) and somatic cells (S-genome) of *E. burgeri* using next-generation sequencing (NGS). Unfortunately, the assembled G-genome contained none of the highly repetitive eliminated families mentioned above, because the copy numbers of these tandem repeats were significantly larger than the average size of the constructed contigs, making their individual assembly impossible. Therefore, an additional K-mer-based analysis was performed to confirm whether the eliminated DNA families found in the previous analysis were also present in the paired-end reads from the germ cells. As a result, not only were 11 eliminated repetitive families reconstructed, but more than 130 additional repetitive DNA families were also identified. These families were reconstructed from read sequences that exhibited frequencies over 100 times higher in the germline reads than in the somatic reads (Goto *et al*., in preparation).

In this study, we first examined the genomic organization and sequence characteristics of four novel and prominent repetitive DNA families. This examination involved PCR amplification and molecular cloning techniques, supported by bioinformatic analysis. Next, to verify the hypothesis of E-chromosome evolution mentioned above, we investigated the chromosomal localization of these four sequences using two-color FISH analysis combined with EEEb1, which is localized to all E- chromosomes in germ cells [8]. Since NGS analysis also showed that four novel repetitive DNA sequences and six other DNA families (EEEb1–6) accounted for about 47.5% and 45.4% of the E- genome, respectively, we inferred the major essence and pattern of E-chromosome evolution by using them for cytogenetical analysis. Hence, to further explore the chromosomal substructure of E- chromosomes, we also mapped these 10 DNA families on metaphase chromosomes and extended chromatin fibers using three-color FISH analysis with confocal microscopy.

## Materials and methods

### Ethics statement

All animal experiments in this study were approved (Protocol#21-52-446) and conducted according to the guidelines of the Institutional Animal Care and Use Committee of Toho University.

### Animals

Japanese hagfish, *Eptatretus burgeri,* were collected from Sagami Bay in Kanagawa, Japan. The animals were injected intraperitoneally with colchicine (0.375 mg/kg body weight) 2 hours before sacrifice to enrich mitotic cells. All animals were euthanized with a high dose of ethyl m-aminobenzoate methanesulfonate (MS-222) (Nacalai Tesque).

### PCR amplification and molecular cloning

We selected four of the novel repetitive DNA families according to their copy number or repeat unit length in descending order. To perform PCR amplification on these selected families (family 0, family 38, family 5, and family 10), we extracted genomic DNA from both germline and somatic tissues (specifically testes and livers). The extraction process followed a standard protocol involving proteinase K treatment, phenol/chloroform extraction, and ethanol precipitation, as previously described [6].

Primer pairs for the amplification of four novel repetitive DNA families were designed by Primer3plus (http://www.bioinformatics.nl/cgi-bin/primer3plus/primer3plus.cgi) with the predicted sequence of each repetitive DNA family and are summarized in Table 1. Each DNA family was amplified with 2×GoTaq^®^ Hot Start Green Master Mix (Promega) using 0.8–2.2 ng of germline and somatic DNA and the primer pair according to the manufacturer’s instructions. Amplification was performed on a T100TM thermal cycler (Bio-Rad Laboratories) under the conditions listed in Table 2.

**Table 1.**
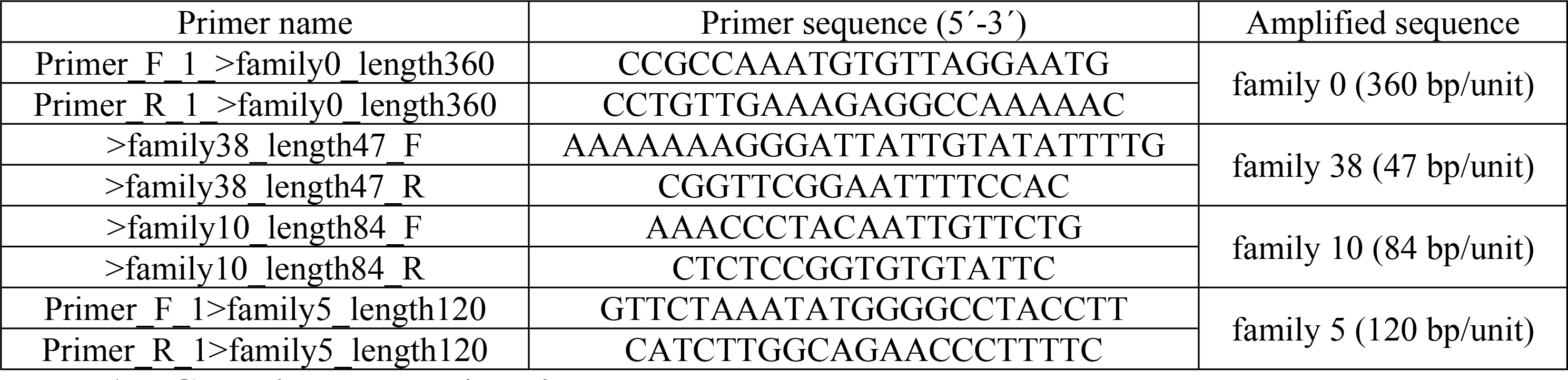
PCR primers used in this study.

**Table 2.** Conditions of the PCR amplification.

| Amplified sequence | Preheat | 35 Cycles |  |  | Final extension |
| --- | --- | --- | --- | --- | --- |
|  |  | Denaturation | Annealing | Extension |  |
| family 0 | 95°C, 2 min | 95°C, 30 sec | 53°C, 30 sec | 72°C, 1 min | 72°C, 5 min |
| family 38 | 95°C, 2 min | 95°C, 30 sec | 57°C, 30 sec | 72°C, 30 sec | 72°C, 5 min |
| family 10 | 95°C, 2 min | 95°C, 30 sec | 55°C, 30 sec | 72°C, 15 sec | 72°C, 5 min |
| family 5 | 95°C, 2 min | 95°C, 30 sec | 57°C, 30 sec | 72°C, 30 sec | 72°C, 5 min |

The PCR products were separated on 2.0% agarose gel for family 0 and 3.0% agarose gel for the other families with a molecular size marker (Gene Ladder wide1; Nippon Gene). The PCR products were purified and ligated into the pMD-20 vector or the pANT vector using the Mighty TA-cloning Kit (Takara) or TA-Enhancer Cloning Kit (Nippon Gene) according to the protocols recommended by the respective suppliers. After transformation into DH5α competent cells (Toyobo) and subsequent blue/white selection on ampicillin plates, plasmid DNAs were prepared from positive clones, and the inserted DNA was sequenced as previously described [25].

### Sequence analysis

All nucleotide sequences were aligned by GENETYX-MAC ver. 19.0.2, which was manually modified when necessary. Gap sites were not included in the calculation of intraspecific sequence diversity. Homology searches were accomplished by using the DNA databases blastn-NCBI (https://blast.ncbi.nlm.nih.gov/Blast.cgi) and Repbase-GIRI (https://www.girinst.org/repbase/).

### FISH

Chromosome preparations from testes and gills were prepared according to the protocol by Nagao *et al*. [11]. Subsequently, these slides were treated with 100 μg/mL RNase A (type I-AS; Merck) in 1xSSC for 30 min at 37°C, followed by dehydration and drying through 70% and 100% ethanol series.

For two-color FISH, the plasmid DNAs, namely F0G_No7, F38G_No2, F10G_No20, and F5G_No5, were labeled with biotin-16-dUTP (Promo Kine), while plasmid DNA containing previously cloned EEEb1 [8], was labeled with digoxigenin-11-dUTP (ENZO) by nick translation as described by Green *et al*. [26]. The labeled probe DNA was then precipitated with 25 μg of yeast tRNA (Invitrogen) and resuspended in 20 μL formamide. After denaturation at 75°C for 10 min, hybridization, washing, and detection were carried out as previously described [11]. Digoxigenin-labeled probes and biotin-labeled probes were detected with 4 µg/mL of anti-digoxigenin Fab fragments conjugated with FITC (Merck) and 1/1,000 diluted streptavidin conjugated with DyLight 549 (Vector Laboratories) in TBST. The slides were counterstained with 0.4 μg/mL Hoechst 33342 (Thermo Fisher Scientific) in TBST. Immunofluorescence and DNA FISH images were captured using an Axio Imager.A2 microscope (Carl Zeiss) equipped with a CCD camera (Carl Zeiss) and Axio vision (Carl Zeiss).

For three-color FISH, plasmid DNA containing EEEb1 was labeled with cyanine-5-dUTP by Nick Translation Mix (Roche), while plasmid DNA harboring each of the other nine DNA families (EEEb2–EEEb10) was labeled with digoxigenin-11-dUTP using DIG-Nick Translation Mix (Roche) or with biotin-16-dUTP (Promo Kine) using Nick Translation Mix (Roche). The plasmid DNA containing EEEb2, EEEb3, EEEb4, EEEb5, or EEEb6, which were previously cloned [11], was used in this experiment. Denaturation of chromosomal DNA and hybridization were performed as described above. After hybridization, the slides were washed in 50% formamide/2xSSC for 15 min at 37°C, 2xSSC for 30 min, and TNT for 5 min at room temperature. After pretreatment with 1/2- diluted Blocking One Histo (Nacalai Tesque) in double-distilled water for 10 min at room temperature, the slides were incubated with 4 µg/mL of anti-digoxigenin Fab fragments conjugated with FITC (Merck) and 1/2,000-diluted streptavidin conjugated with alexa405 (Vector Laboratories) or 1/2,000- diluted streptavidin conjugated with cyanine-3 (Vector Laboratories) in TNT for 1 hour at room temperature in a dark humid chamber. After the slides were washed three times with TNT and counterstaining with 0.05 μg/mL propidium iodide (Fujifilm Wako) or 0.4 μg/mL Hoechst 33342 in TNT, they were mounted with Fluoro-KEEPER Antifade Reagent (Nacalai Tesque). Immunofluorescence images and DNA FISH images were captured using a Nikon A1R confocal microscope. The microscope was controlled by Nikon NIS-Elements software. A Plan Apo ×100/1.4 NA oil immersion objective was used for imaging. Lasers with excitation lines at 405, 4880, 594, and 633 nm were employed for fluorescence excitation during image acquisition. The fluorescence signal distribution was analyzed using ImageJ2 (Fiji) ver. 2.3.0/1.53q.

### Fiber-FISH

The testis was homogenized in a microcentrifuge tube containing hagfish ringer’s solution (0.5 M NaCl, 8 mM KCl, 2 mM CaCl_2_, 2 mM NaHCO_3_, 4 mM MgCl_2_) on ice. The tube was then briefly centrifuged at 1500 rpm for a few seconds at room temperature, and the supernatant was aspirated. The cell pellet was then resuspended in a 2/3-diluted hagfish ringer’s solution and subjected to centrifugation under the same conditions. This process was repeated several times, after which the suspensions were stored at 4°C until chromatin preparation. Thereafter, 15 μL of the cell suspension was spread on glass slides in a circular pattern and hemi-dried at room temperature before crystallization of the droplet edge. The slides were vertically immersed in lysis solution (25 mM Tris-HCl [pH 8.0], 1% TritonX-100, 2 M NaCl) for 5 seconds at room temperature and slowly removed from the lysis solution. The preparation was immediately fixed by immersion in 70% ethanol for 30 min at room temperature. The slides were then immersed in 100% ethanol for 5 min, air dried, and stored at -20°C until the following experiments. Probe labeling, RNase treatment, denaturation of chromosomal DNA, hybridization, washing, detection, image capture, and image analysis were carried out as described above for three-color FISH analysis.

## Results

### PCR amplification and molecular cloning of novel repetitive DNA families

To investigate the genomic organization and sequence characteristics of the newly discovered sequences, four of them, namely families 0, 38, 10, and 5 (sorted by their copy numbers), were amplified by PCR using germline and somatic DNA with each primer pair summarized in Table 2 and separated on agarose gel (Fig 1a). In family 0, both 360 bp and 720 bp DNA bands were amplified from somatic DNA, suggesting the specific amplification of the monomer and dimer of family 0. In contrast, not only the same DNA bands but also additional DNA bands corresponding to 450, 900, and 1100 bp were detected in PCR using germline DNA. In the cases of family 38 (47 bp/unit) and family 10 (84 bp/unit), the monomers and oligomers of each family were distinctly detected in somatic DNA, while the same bands and longer multimers were observed in smears in the germline DNA lanes. DNA equivalent to the monomer and multimers of family 5 (120 bp/unit) was amplified in both germline and somatic DNA, although highly multimeric repeat DNA was preferentially observed in the germline DNA. These results indicate that all four novel repetitive DNA sequences were tandemly repeated in the germline and somatic genomes and were multimerized more extensively in the germline genome than in the somatic genome.

**Fig 1.**
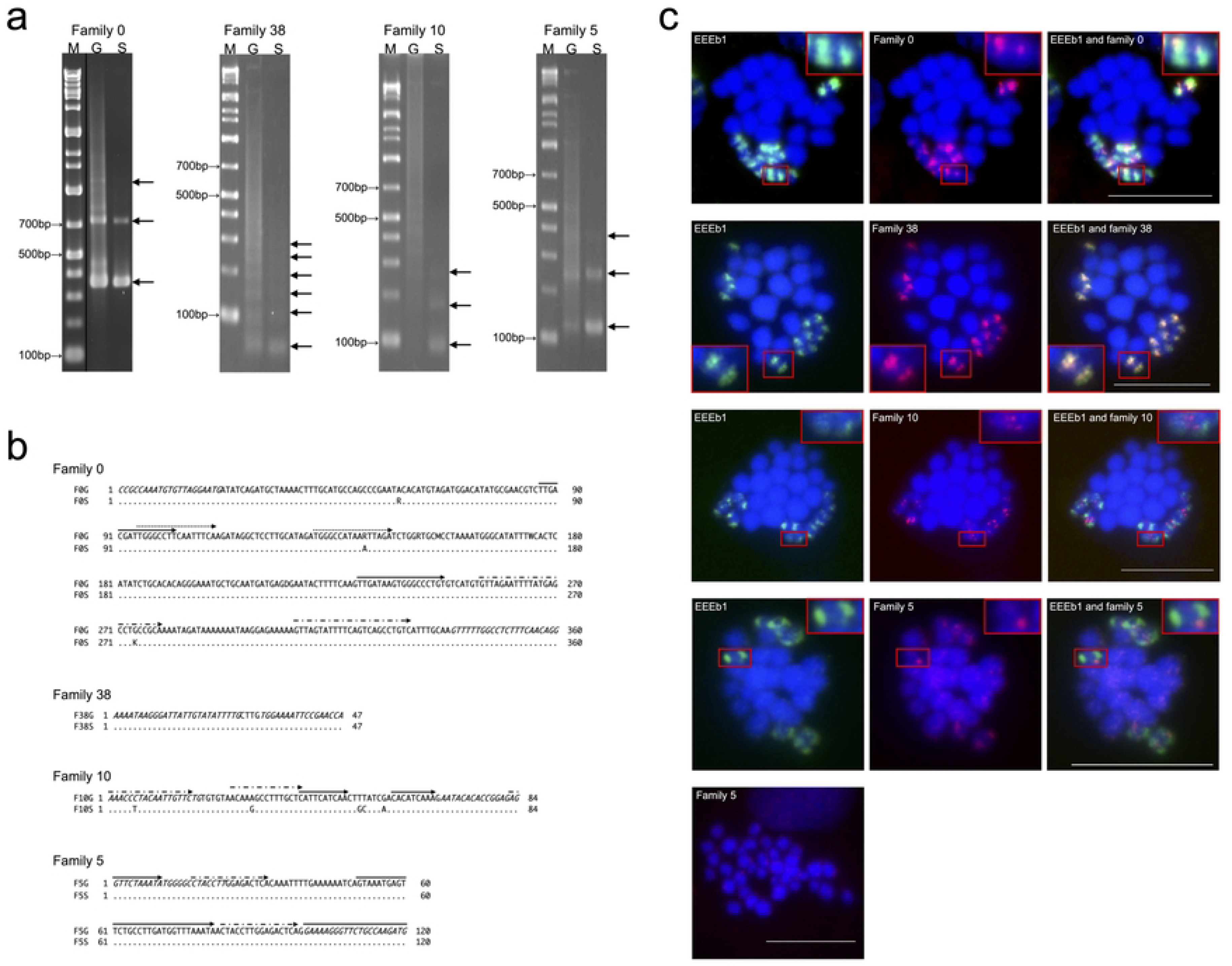
Identification of four novel eliminated repetitive DNA families. **(a)** Amplification of four novel repetitive DNA families. PCR products using germline DNA (lane G) and somatic DNA (lane S) of *E. burgeri* as templates were separated on 3.0% agarose gel (family 0) or 2.0% agarose gel (families 38, 10, and 5). The left lane in the image contains the DNA molecular size marker. The corresponding sizes of the monomers and multimers of each family are indicated by arrows. (**b**) The consensus sequences of the four repetitive DNA families. Each consensus nucleotide sequence was derived from the inserts of germline DNA clones and somatic DNA clones of family 0 (F0G and F0S), family 38 (F38G and F38S), family 10 (F10G and F10S), and family 5 (F5G and F5S). Germline sequences (top) and somatic sequences (bottom) are aligned. Nucleotides identical to those in the consensus sequence at the top are represented by dots (.), while base substitutions are indicated by the respective bases. Primer regions are marked with italic letters and direct repeats by arrows. The obtained sequence data were deposited in GenBank (LC731298–LC731305). (**c**) Chromosomal mapping of the four DNA families in *E. burgeri*. The digoxigenin-labeled EEEb1 probes (green) and biotin-labeled families 0, 38, 10, and 5 (red) were hybridized on the metaphase chromosomes in the spermatocytes. The chromosomal localization of family 5 in somatic cells is also shown at the bottom panel. Chromosomes were counterstained with Hoechst 33342 (blue). The magnified images of E-chromosomes with each signal are shown in insets. The scale bar was set to 5 μm.

All novel DNA families amplified by PCR from both germline and somatic DNAs were independently cloned and sequenced. The resulting consensus sequences for families 0, 38, 10, and 5 were determined from 16, 40, 34, and 12 examined sequences, respectively, obtained from the germline genome. These sequences, with repeat unit lengths of 360, 47, 84, and 120 bp were respectively designated F0G, F38G, F10G, and F5G (Fig 1b; top). Similarly, four consensus sequences were deduced from 10, 50, 21, and 22 somatic genome-derived sequences, designated F0S, F38S, F10S, and F5S, respectively, and were aligned with germline-derived consensus sequences to determine their homology (Fig 1b; bottom). Comparison of germline and somatic genome-derived consensus sequences showed three and five nucleotide substitutions in families 0 and 10, respectively, with GC contents of 40.8% (F0G), 40.9% (F0S), 39.3% (F10G), and 40.5% (F10S). In contrast, no base substitutions were detected in families 5 and 38, with GC contents of 29.8% and 39.2%, respectively. The average intraspecific divergence of the four repetitive families in the germline and somatic genomes were 4.5% and 5.3% in family 0, 7.4% and 7.1% in family 38, 18.6% and 11.0% in family 10, and 2.2% and 0.4% in family 5. Three, two, and two pairs of direct repeats were present in the consensus sequences of families 0, 10, and 5, respectively (Fig 1b; arrows), with no continuous open reading frames (ORF) detected in any of the four families. Family 5, however, showed homology to the vicinity of Hox genes in *E. burgeri* (GenBank accession numbers MF182102 and MF182105; Pascual-Anaya *et al*. [27]), with three copies of family 5 found in tandem in both MF182102 and MF182105. No significant homologies were found in the other consensus sequences.

### Chromosomal mapping of repetitive DNA families

To ascertain whether the four DNA families belonged to the E-genome, their chromosomal localization was examined through two-color FISH analysis. EEEb1, which is exclusively distributed on all E-chromosomes, was simultaneously detected and used as a control (Fig 1c). In our previous study, we hypothesized that the eight pairs of E-chromosomes are derived from one pair of ancestral chromosomes with multiple duplication events during a long evolutionary period, as the other six DNA families were located on all E-chromosomes with similar distributions [11]. If this hypothesis is true, it is expected that the novel eliminated DNA families would also be located on all E- chromosomes. In family 0, fluorescent signals were detected on all EEEb1-positive chromosomes but not on any EEEb1-negative chromosomes in metaphases of spermatocytes. Image analysis revealed that family 0 signals were mostly incorporated into those of EEEb1, although they were partly detected in the vicinity of EEEb1 signals. Family 38 also appeared to cluster on all EEEb1-positive chromosomes in the first meiotic metaphase, with EEEb1-negative chromosomes having no family 38 signals. The distribution and size of family 38 signals were completely consistent with those of EEEb1. In the case of family 10, significant signals were primarily detected in regions neighboring the EEEb1 signals. Furthermore, the minor signals exhibited co-localization with the EEEb1 signals on EEEb1-positive chromosomes within metaphase spermatocytes. No family 10 signals were ever observed on EEEb1-negative chromosomes. Family 5 signals were weakly detected on all chromosomes in germ cells as scattered signals, and the signals detected on EEEb1-positive chromosomes tended to be intense. Weak signals were detected on all chromosomes in somatic cells. These four families, predicted to be E-genome by K-mer-based analysis, were all distributed on the eliminated chromosomes and were designated EEEb7, EEEb8, EEEb9, and EEEb10. These findings support the aforementioned hypothesis regarding E-chromosome evolution [11].

In addition to these four families, an additional six repetitive families were previously identified as E-genome [11]. Since these 10 DNA families account for >90% of the E-genome, we can infer the evolutionary pattern of E-chromosomes and verify the previously established hypothesis [11] more precisely. Therefore, to investigate the relative positions of the 10 families, we performed three-color FISH combining two of the aforementioned families with EEEb1 in spermatocytes (Fig 2). Our findings were consistent with previous studies [8, 11] and our present study (Fig 1c), wherein fluorescent signals for each DNA family were detected on all EEEb1-positive chromosomes in the metaphase of spermatocytes. Using image analysis, we divided the signal distributions of the nine DNA families (EEEb2–10) on E-chromosomes into four groups by comparing them with the distributions of EEEb1, despite the similarity in their individual E-chromosome distributions. Specifically, (*i*) the signal clusters of EEEb8 were fully aligned with EEEb1 clusters; (*ii*) EEEb2, EEEb4, and EEEb7 largely overlapped with EEEb1 clusters; (*iii*) half of the signals of EEEb6 and EEEb9 were overlapped with EEEb1 clusters; and (*iv*) those of EEEb3, EEEb5, and EEEb10 were predominantly located on EEEb1-negative regions and the peripheries of EEEb1-positive regions.

**Fig 2.**
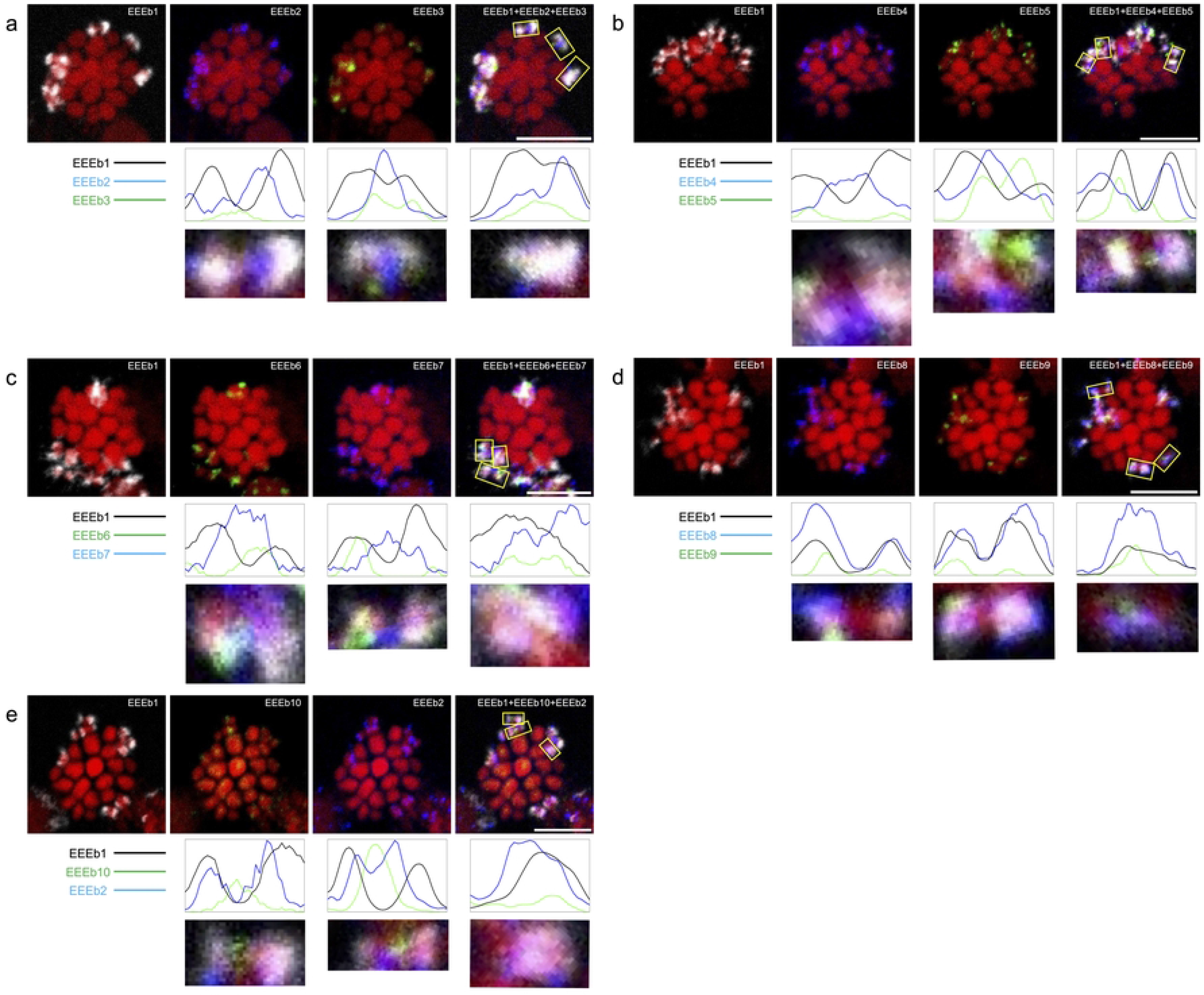
Chromosomal mapping of 10 eliminated DNA families in *E. burgeri*. Metaphase chromosomes in spermatocyte were hybridized using cyanine-5-labeled EEEb1 probes (gray) along with biotin-labeled EEEb2 (**a** and **e**), EEEb4 (**b**), EEEb7 (**c**), and EEEb8 (**d**) (blue) as well as digoxigenin-labeled EEEb3 (**a**), EEEb5 (**b**), EEEb6 (**c**), EEEb9 (**d**), and EEEb10 (**e**) (green) (top). Fluorescence intensity histograms, derived from XY-sections, were displayed below representative individual E-chromosomes (bottom). Chromosomes were counterstained with propidium iodide (red). The scale bar was set to 5 μm.

Based on this categorization, additional FISH experiments were conducted using combinations of each DNA family classified into the same signal distribution group as the EEEb1 probe (Fig 3). As shown in Fig 3a–3c, the signal localizations of EEEb2, EEEb4, and EEEb7 were slightly different, although all were classified into the same signal distribution group. Half of the signals overlapped with each other on the EEEb1-positive regions of all E-chromosomes, while the remaining signals were exclusively detected on the EEEb1-negative regions of certain E-chromosomes. On the other hand, the signals of EEEb6 predominantly corresponded to those of EEEb9 on both the EEEb1-negative and -positive regions of all E-chromosomes (Fig 3d). In the cases of EEEb3 and EEEb5, their signals exhibited scarce concordance on both the EEEb1-positive and - negative regions of some of the E-chromosomes (Fig 3e). Some of the EEEb5 signals overlapped with the intense EEEb10 signals on both the EEEb1-positive and -negative regions of the E- chromosomes (Fig 3f). The EEEb3 signals tended to coincide with the intense EEEb10 signals on both the EEEb1-positive and -negative regions of E-chromosomes (Fig 3g).

**Fig 3.**
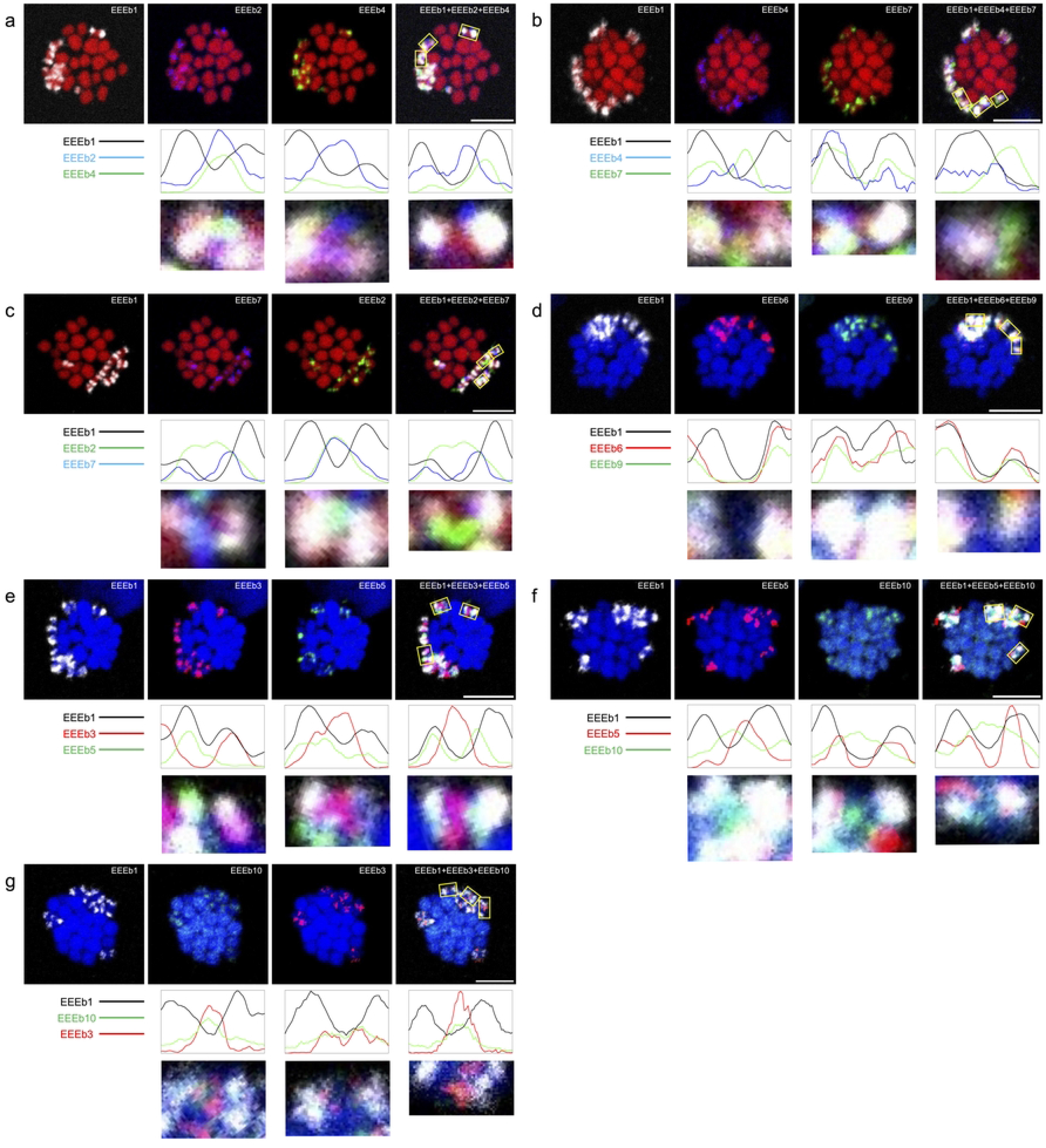
Chromosomal mapping of similarly distributed DNA families in *E. burgeri*. Metaphase chromosomes in spermatocytes were hybridized using cyanine-5-labeled EEEb1 probes (gray) in combination with biotin-labeled EEEb2 (blue) and digoxigenin-labeled EEEb4 (green) (**a**), digoxigenin-labeled EEEb4 (green), and biotin-labeled EEEb7 (blue) (**b**), biotin-labeled EEEb7 (blue) and digoxigenin-labeled EEEb2 (green) (**c**), biotin-labeled EEEb6 (red) and digoxigenin-labeled EEEb9 (green) (**d**), biotin-labeled EEEb3 (red) and digoxigenin-labeled EEEb5 (green) (**e**), biotin-labeled EEEb5 (red) and digoxigenin-labeled EEEb10 (green) (**f**), or digoxigenin-labeled EEEb10 (green) and biotin-labeled EEEb3 (red) (**g**). Chromosomes were counterstained with propidium iodide (**a**–**c**) (red) or Hoechst33342 (**d**–**g**) (blue). The other notations correspond to those in Fig 2.

Using these hybridization signals, we constructed a karyogram encompassing eight E- chromosomes, along with a map illustrating the locations of EEEb1–10 DNA families on individual E-chromosomes (Fig 4). Further examination revealed that the eight pairs of E-chromosomes could be classified into six distinct patterns (patterns 1–6) based on the distribution of signal intensity for each DNA family. Notably, two pairs of E-chromosomes exhibited identical signal distribution patterns (patterns 1 and 2). In EEEb1 and EEEb8, the signal localization patterns exhibited a dichotomy: (*i*) two clusters were symmetrically located on the terminal regions of seven pairs of E- chromosomes (patterns 1–5), and (*ii*) a single cluster was located on the terminal region of one pair of E-chromosomes (pattern 6). Conversely, the chromosomal localization patterns of the remaining eight DNA families showed intricate and distinct distribution patterns. However, in most cases, the signal localization of each DNA family exhibited an almost symmetrical distribution across all E- chromosomes except for pattern 6 E-chromosomes. Patterns 1–5 E-chromosomes exhibited symmetrical distribution patterns for at least seven of the eliminated DNA families, while pattern 6 E-chromosomes lacked symmetrical signal distribution for any DNA families. This remarkable symmetry observed in the karyogram strongly suggests an isochromosomal nature for patterns 1–5 E-chromosomes in this species.

**Fig 4.**
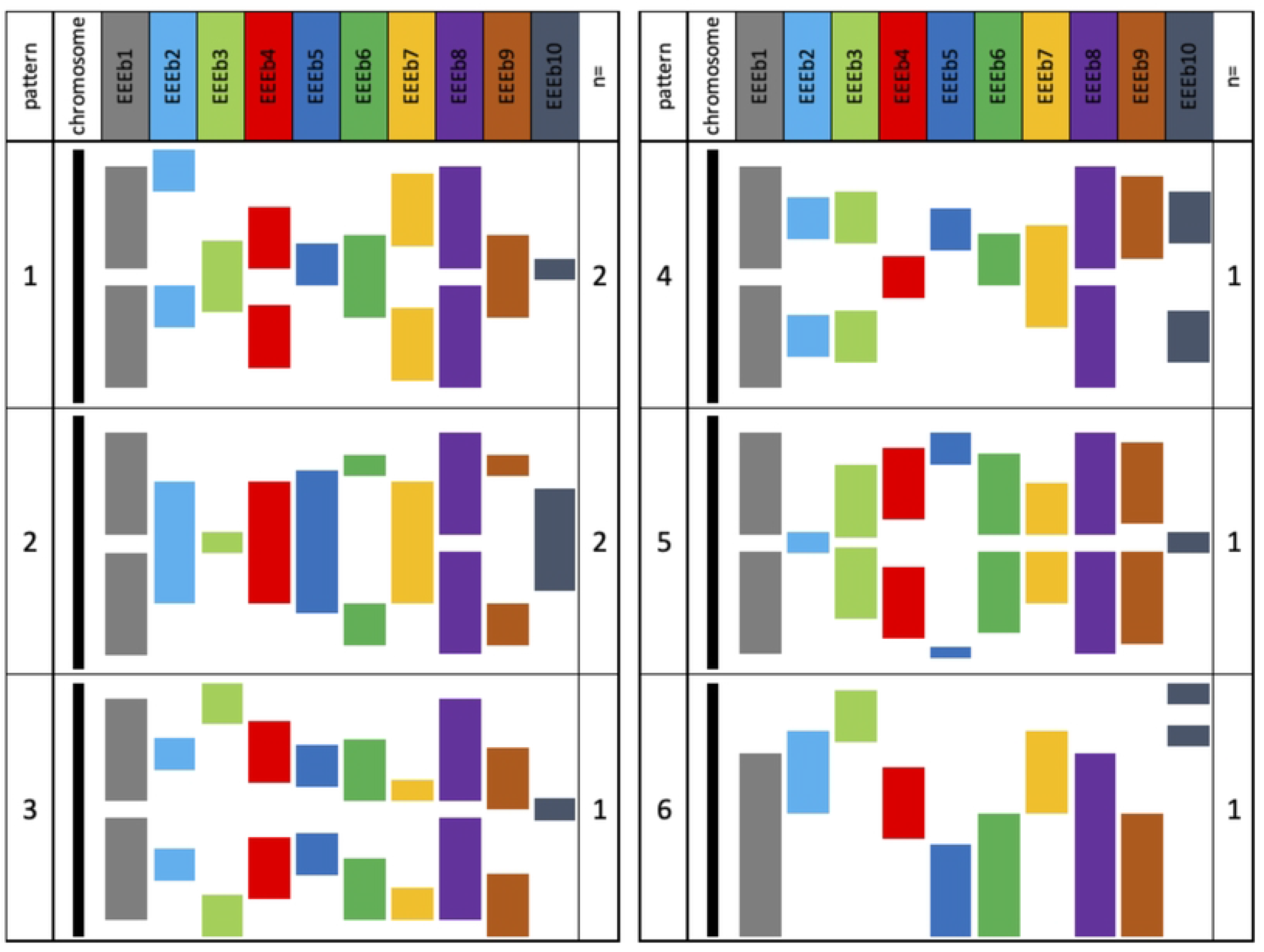
A schematic karyogram of distribution patterns of 10 eliminated DNA families in *E. burgeri*. The karyogram is constructed based on the image analysis of FISH images using EEEb1 to EEEb10 probes (Figs 2 and 3). In the case of EEEb10, although weak signals were detected across the entire regions of E-chromosomes and somatically retained chromosomes, the karyogram selectively emphasized the regions exhibiting intense signals. Eight pairs of E-chromosomes were divided into six patterns according to the signal distribution patterns of EEEb1–EEEb10 in metaphase chromosomes of spermatocytes (patterns 1 to 6). E-chromosomes belonging to patterns 1–5 showed symmetric distribution patterns across at least seven eliminated DNA families, while E-chromosomes in pattern 6 did not exhibit symmetric signal distribution for any DNA families. The notation n= indicates chromosomal pair(s) of the E-chromosomes categorized into each pattern.

### Fine mapping of eliminated repetitive DNA families

To gain a more detailed understanding of the distribution of EEEb1–10 within chromosome E, fiber-FISH analysis was performed on elongated chromatin fibers obtained from testicular cells, using the same probe combination as in Figure 2. Within this experiment, the EEEb1 probe served as a marker to specifically identify chromatin fibers derived from E- chromosomes on meiotic metaphase chromosomes. As shown in Fig 5, the fiber-FISH analysis revealed the interspersed distribution of EEEb1 signals along the extended chromatin fibers (Fig 5a–5e; top panels). Similarly, the signals from the other nine eliminated DNA families (EEEb2 to EEEb10) were also dispersed within EEEb1-positive chromatin fibers, appearing intertwined with both EEEb1 signals and with each other. This was observed through multi-color FISH analysis conducted on metaphase chromosomes in spermatocytes.

**Fig 5.**
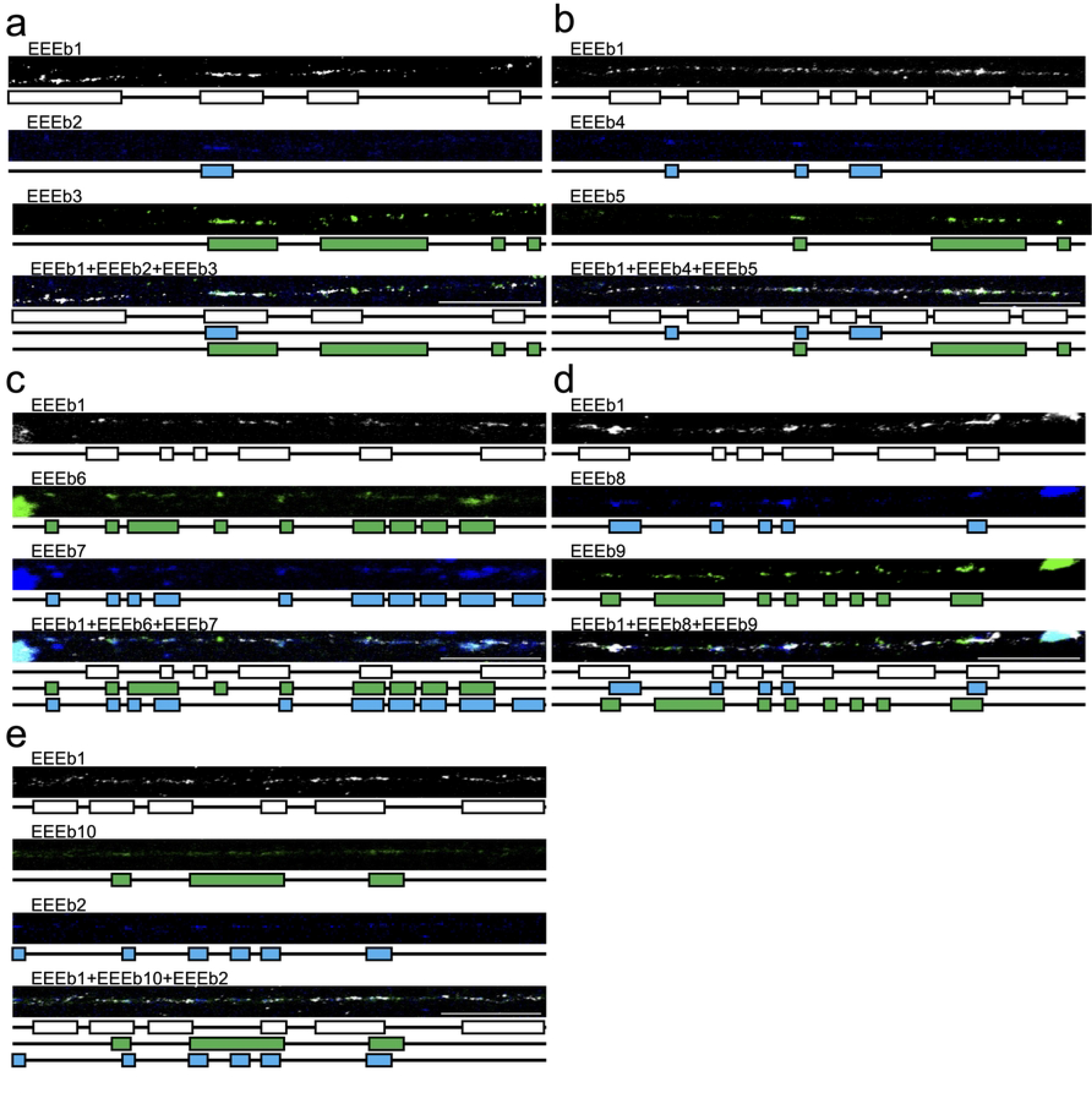
Fiber-FISH analysis of 10 eliminated DNA families in *E. burgeri*. Extended chromatin fibers derived from testicular cells were subjected to hybridization using cyanine-5-labeled EEEb1 probes (gray) with biotin-labeled EEEb2 (**a** and **e**), EEEb4 (**b**), EEEb7 (**c**), and EEEb8 (**d**) (blue). Additionally, digoxigenin-labeled EEEb3 (**a**), EEEb5 (**b**), EEEb6 (**c**), EEEb9 (**d**), and EEEb10 (**e**) were also utilized for the hybridization and were detected (green). Schematic diagrams corresponding to each signal image appear below. Scale bar=20 µm.

Based on the signal distribution of each DNA family on metaphase chromosomes, additional fiber-FISH experiments were conducted using the same probe combination as in Fig 3 (Fig 6). Consistent with the findings from the multicolor FISH on metaphase chromosomes, the signals of the nine eliminated DNA families (EEEb2 to 10) were interspersed along EEEb1-positive chromatin fibers, entwined with EEEb1 signals and with one another.

**Fig 6.**
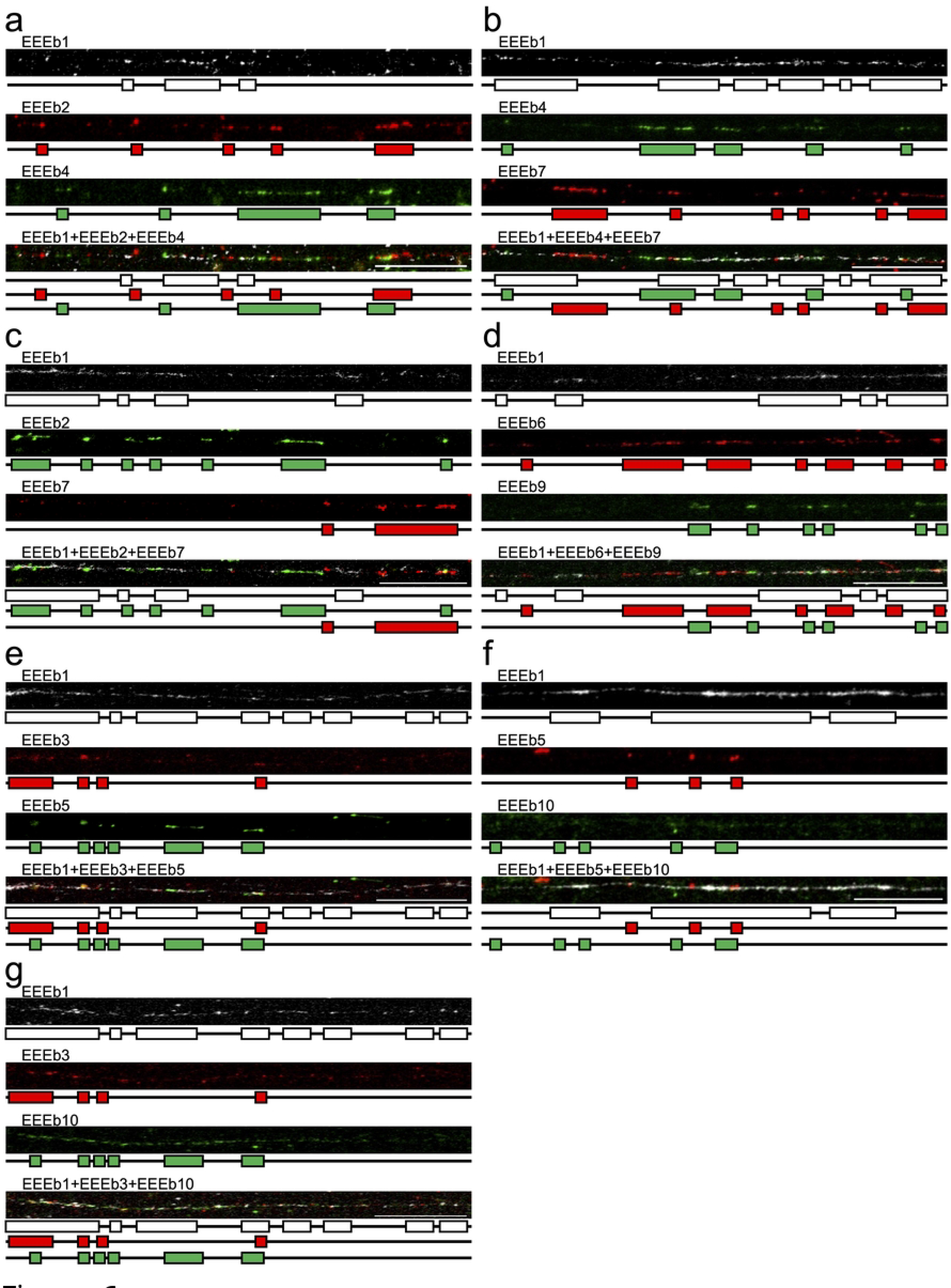
Fiber-FISH analysis of similarly distributed DNA families in *E. burgeri*. Extended chromatin fibers from testicular cells were hybridized using cyanine-5-labeled EEEb1 probes (gray) with biotin-labeled EEEb2 (red) and digoxigenin-labeled EEEb4 (green) (**a**), digoxigenin-labeled EEEb4 (green) and biotin-labeled EEEb7 (red) (**b**), digoxigenin-labeled EEEb2 (green) and biotin-labeled EEEb7 (red) (**c**), biotin-labeled EEEb6 (red) and digoxigenin-labeled EEEb9 (green) (**d**), biotin-labeled EEEb3 (red) and digoxigenin-labeled EEEb5 (green) (**e**), biotin-labeled EEEb5 (red) and digoxigenin-labeled EEEb10 (green) (**e**), or biotin-labeled EEEb3 (red) and digoxigenin-labeled EEEb10 (green) (**g**). The other notations correspond to those in Fig 5.

## Discussion

In this study, four novel repetitive DNA families were identified as the eliminated repetitive sequences of *E. burgeri*. These families were predicted through a bioinformatics approach utilizing NGS data. The identification process involved molecular genetic techniques such as PCR amplification, as well as cytogenetic methods including two-color FISH analysis. Our investigations clearly demonstrated that all of the families that were predicted to account for 47.5% of their E-genome based on NGS analysis (that is, EEEb7 to EEEb10) are tandemly repeated and specifically localized on E-chromosomes. Furthermore, three-color FISH analysis using a combination of 10 repetitive DNA families (EEEb1–10), which constitute approximately 90% of the E-genome, revealed that all E-chromosomes could be classified into six groups based on the distribution patterns of the signals. Additionally, fiber-FISH analysis with these 10 eliminated DNA families also showed the intermingled signal distribution with each other on extended chromatin fibers.

It is well established that E-chromosomes in animals undergoing PGR, including hagfish species, consist predominantly of heterochromatic regions and encompass numerous repetitive DNA families [6-10, 28-31]. Despite extensive investigations conducted to elucidate the evolutionary mechanisms involved in chromosome elimination, the evolutionary origin or origins of E- chromosomes in most species remain unknown. In the case of chironomids and songbirds, which eliminate germline-restricted chromosomes (GRCs) during early embryogenesis [32, 33], microdissected GRC-specific probes were clearly detected not only on the GRC itself but also on a pair of autosomal chromosomes that are retained in the somatic cells. This finding strongly indicates that the GRCs originated from the intraspecific complement of autosomes [30, 34, 35]. On the other hand, in the case of the gnat, which also undergoes PGR during early embryogenesis, different results have been reported from comparative genome analysis between related lineages. In the Sciaridae gnat, *Bradysia coprophila*, the majority of genes found on the GRCs exhibited significant divergence from their paralogs on the autosomes, while these GRC genes showed sequence similarity to the orthologs present in the Cecidomyiidae gnat lineage. These results suggest that the GRCs of the Sciaridae gnat likely arose through introgression from the Cecidomyiidae lineage via a hybridization event during the early divergence between Sciaridae and Cecidomyiidae [22, 36]. In contrast to the Sciaridae gnat, the E-chromosome of *E. burgeri*, particularly EEEb1, contains components that have not been detected in other hagfish species [8]. Additionally, EEEb10 is shared among all E-chromosomes and the remaining chromosomes retained in somatic cells (Fig 1c). These observations suggest that the E-chromosomes of hagfish species, at least in the case of *E. burgeri*, likely originated from intraspecific autosomes (somatically retained chromosomes). This origin involves the loss and replacement of sequences from one or more ancestral autosomes by eliminated repetitive DNA families, such as EEEb1 and EEEb7, rather than chromosomes originating from different species.

In our previous study, FISH analysis revealed a consistent signal distribution of EEEb1 to EEEb6 on all eight pairs of E-chromosomes [11]. Based on this specific detection, we hypothesized that the eight pairs of E-chromosomes originate from a single pair of ancestral chromosomes through multiple duplication events. These events were likely driven by meiotic drive, leading to copy number variations and chromosomal rearrangements over a prolonged evolutionary period. To further support this hypothesis, the present study showed that EEEb7 to EEEb9, but not EEEb10, were mapped on all E-chromosomes (Fig 1c). These findings suggest that, by using other predicted repetitive DNA families as FISH probes, visible signals would exclusively localize to all E-chromosomes of *E. burgeri* as EEEb1–EEEb9. Similar results were reported in *E. okinoseanus*, where two eliminated sequences, EEEo1 and EEEo2, were extensively observed on all E-chromosomes [6]. However, in other examined hagfish species, such as *E. cirrhatus* and *P. sheni*, the repetitive sequences that were identified as eliminated repeats were mapped on some, but not all, of the E-chromosomes [9, 10]. These findings suggest that the evolution of E-chromosomes in hagfish species can be categorized into two groups. In one group, represented by *E. burgeri* and *E. okinoseanus*, all E-chromosomes display a high degree of homogeneity and have likely undergone multiple duplication events. This implies a shared evolutionary history and a concerted pattern of evolution for the E-chromosomes in these species. In contrast, the other group, represented by *E. cirrhatus* and *P. sheni*, shows less homogeneity among their E-chromosomes, indicating independent evolutionary trajectories for each individual E-chromosome. Further comparative cytogenetic analysis can provide valuable insights into the evolutionary mechanisms underlying the diversification of E-chromosomes in hagfish species.

In *E. burgeri,* although the chromosomes are morphologically similar, the E-chromosomes could be divided into six groups based on the distribution patterns of the 10 eliminated DNA families during meiotic metaphase-I (Fig 4). The presence of two pairs of chromosomes with pattern 1 and pattern 2 indicates that these chromosome groups have remained conserved without significant chromosomal rearrangements within each group following the chromosome duplication event. Among the examined DNA families, EEEb1 and EEEb8 in particular showed symmetric distributions in patterns 1–5 E-chromosomes, but pattern 6 did not (Fig 4). However, the degree of conservation of symmetric distribution varied among different eliminated DNA families and between groups. In Characidae fish species, supernumerary (B) chromosomes, which are additional dispensable chromosomes that occur frequently among multicellular organisms [37], are known to be present with intraspecific variants [38–41]. Some of these B chromosome variants exhibit isochromosome features, characterized by the symmetrical distribution of repetitive DNA families that are found on intraspecific autosomes. Additionally, meiotic self-pairing between the two arms of the chromosome is observed. These findings indicate that the B chromosome variants with isochromosome characteristics originated from intraspecific autosomes through the incorrect division of the acrocentric chromosomes. Similarly, in *E. burgeri*, although the chromosomal arms are morphologically indistinguishable [4], it is possible that the E-chromosomes originated as isochromosomes through chromatid nondisjunction and chromosomal fusion events involving ancestral chromosomes, similar to what is observed in Characidae species.

To date, FISH analyses using 12 eliminated DNA families (EEEb1–10, EEEo1, and EEEo2) have revealed that all of the eliminated DNA families were exclusively detected on either E- chromosomes or somatically retained chromosomes in *E. burgeri*, with the exception of EEEb10, which was detected on both germline-restricted and somatically retained chromosomes (Fig 1c) [8, 11, 25]. Although the genomic contents were conserved among E-chromosomes, indicating the presence of an autonomous mechanism such as recombination-mediated DNA repair system to maintain the repetitive sequence array, these findings imply limited homogenization of the genomic contents between the germline-restricted and somatically retained chromosomes in this species. In bird species, their karyotypes usually have large numbers of diploid chromosomes, typically around 80, and consist of morphologically distinguishable macrochromosomes and microchromosomes. Centromeric repetitive sequences are preferentially amplified in either several macrochromosomes or all microchromosomes [42–45]. These findings indicate that centromeric repetitive sequences are homogenized within macrochromosomes or microchromosomes, with the homogenization process being more pronounced among microchromosomes. In chicken interphase nuclei, macrochromosomes and microchromosomes show different spatial organizations: the former cluster in the nuclear center while the latter are distributed at the nuclear periphery [46, 47]. Considering these findings, it is evident that DNA transposition (homogenization) through unequal crossing over frequently takes place in the centromeric regions between physically close, nonhomologous macro- or microchromosomes. Although no specific chromosomal organization has ever been observed in the interphase nuclei of *E. burgeri*, the bivalent E-chromosomes tended to assemble each other in the mitotic metaphase nuclei [4]. This may explain the differential conservation of genomic contents between E-chromosomes and somatically retained chromosomes in this species.

Fiber-FISH analysis of extended chromatin fibers from germ cells of *E. burgeri* revealed that the signal localizations of the 10 eliminated repetitive DNA families were intermingled with each other, indicating the absence of clear boundaries between clusters of different DNA families at the chromatin level (Fig 5). To the best of our knowledge, fiber-FISH analysis on animals exhibiting PGR has not been previously conducted. However, the intermingled arrangement of nonhomologous repetitive DNA families has been observed in diverse taxa [48–55]. For instance, in the grasshopper species *Abracris flavolineata*, FISH analysis showed that two distinct DNA families, namely AflaSAT-1 and AflaSAT-2, were colocalized on the large pericentromeric region of all autosomal chromosomes and the small centromeric region of B chromosomes. Subsequent fiber-FISH analysis also demonstrated that these two DNA families were intermingled with each other on the extended chromatin fibers [53]. These findings indicate that, although tandemly repetitive DNA families appear as a single visible FISH signal cluster on metaphase chromosomes, the clusters of different DNA families are intermixed and nested within each other at the chromatin level, rather than a single DNA family being exclusively repeated in a tandem array. However, the specific boundary sequences between DNA families remain unclear. Hence, further investigation of both the genomic contents and chromosomal organization will provide insights into the evolution of E-chromosomes and PGR.

## Acknowledgments

We are grateful to Mr. S. Hatanaka of Shinshomaru, Katase Coast, for supplying experimental materials. We also thank Tomoyuki Aizu and Hinako Ishizaki at the National Institute of Genetics for supporting the NGS analysis.

## Data availability

Sequence data have been deposited in GenBank (accession nos. LC731298–LC731305).

## References

1. Boveri T. Über Differenzierung der Zellkerne während der Furchung des Eies von *Ascaris megalocephala*. Anat Anz. 1887;2:688–93.

2. Wang J, Davis RE. Programmed DNA elimination in multicellular organisms. Curr Opin Genet Dev. 2014;27:26–34. Epub 20140602. doi: 10.1016/j.gde.2014.03.012. PubMed PMID: 24886889; PubMed Central PMCID: PMCPMC4125452.

3. Dedukh D, Krasikova A. Delete and survive: strategies of programmed genetic material elimination in eukaryotes. Biol Rev Camb Philos Soc. 2022;97(1):195–216. Epub 20210920. doi: 10.1111/brv.12796. PubMed PMID: 34542224; PubMed Central PMCID: PMCPMC9292451.

4. Kohno S, Nakai Y, Satoh S, Yoshida M, Kobayashi H. Chromosome elimination in the Japanese hagfish, *Eptatretus burgeri* (Agnatha, Cyclostomata). Cytogenet Cell Genet. 1986;41(4):209–14. Epub 1986/01/01. doi: 10.1159/000132231. PubMed PMID: 3709235.

5. Kohno S, Kubota S, Nakai Y. Chromatin Diminution and Chromosome Elimination in Hagfishes. In: Jørgensen JM, Lomholt JP, Weber RE, Malte H, editors. The biology of hagfishes. Dordrecht: Springer Netherlands; 1998. p. 81-100.

6. Kubota S, Kuro-o M, Mizuno S, Kohno S. Germ line-restricted, highly repeated DNA sequences and their chromosomal localization in a Japanese hagfish (*Eptatretus okinoseanus*). Chromosoma. 1993;102(3):163–73. Epub 1993/02/01. doi: 10.1007/BF00387731. PubMed PMID: 8458254.

7. Kubota S, Ishibashi T, Kohno S. A germline restricted, highly repetitive DNA sequence in *Paramyxine atami*: an interspecifically conserved, but somatically eliminated, element. Mol Gen Genet. 1997;256(3):252–6. Epub 1997/12/11. doi: 10.1007/s004380050567. PubMed PMID: 9393449.

8. Kubota S, Takano J, Tsuneishi R, Kobayakawa S, Fujikawa N, Nabeyama M, et al. Highly repetitive DNA families restricted to germ cells in a Japanese hagfish (*Eptatretus burgeri*): a hierarchical and mosaic structure in eliminated chromosomes. Genetica. 2001;111(1-3):319–28. Epub 2002/02/14. doi: 10.1023/a:1013751600787. PubMed PMID: 11841177.

9. Goto Y, Kubota S, Kohno S. Highly repetitive DNA sequences that are restricted to the germ line in the hagfish *Eptatretus cirrhatus*: a mosaic of eliminated elements. Chromosoma. 1998;107(1):17–32. Epub 1998/05/06. doi: 10.1007/s004120050278. PubMed PMID: 9567198.

10. Kojima NF, Kojima KK, Kobayakawa S, Higashide N, Hamanaka C, Nitta A, et al. Whole chromosome elimination and chromosome terminus elimination both contribute to somatic differentiation in Taiwanese hagfish *Paramyxine sheni*. Chromosome Res. 2010;18(3):383–400. Epub 20100330. doi: 10.1007/s10577-010-9122-2. PubMed PMID: 20352325.

11. Nagao K, Otsuzumi T, Chinone H, Sasaki T, Yoshimoto J, Matsuda M, et al. Novel selectively amplified DNA sequences in the germline genome of the Japanese hagfish, *Eptatretus burgeri*. Sci Rep. 2022;12(1):21373. Epub 20221209. doi: 10.1038/s41598-022-26007-2. PubMed PMID: 36494570.

12. Nabeyama M, Kubota S, Kohno S. Concerted evolution of a highly repetitive DNA family in eptatretidae (Cyclostomata, agnatha) implies specifically differential homogenization and amplification events in their germ cells. J Mol Evol. 2000;50(2):154–69. Epub 2000/02/23. doi: 10.1007/s002399910017. PubMed PMID: 10684349.

13. Wang J, Mitreva M, Berriman M, Thorne A, Magrini V, Koutsovoulos G, et al. Silencing of germline-expressed genes by DNA elimination in somatic cells. Dev Cell. 2012;23(5):1072–80. Epub 20121101. doi: 10.1016/j.devcel.2012.09.020. PubMed PMID: 23123092; PubMed Central PMCID: PMCPMC3620533.

14. Wang J, Gao S, Mostovoy Y, Kang Y, Zagoskin M, Sun Y, et al. Comparative genome analysis of programmed DNA elimination in nematodes. Genome Res. 2017;27(12):2001–14. Epub 20171108. doi: 10.1101/gr.225730.117. PubMed PMID: 29118011; PubMed Central PMCID: PMCPMC5741062.

15. Wang J, Veronezi GMB, Kang Y, Zagoskin M, O’Toole ET, Davis RE. Comprehensive chromosome end remodeling during programmed DNA elimination. Curr Biol. 2020;30(17):3397-413 e4. Epub 20200716. doi: 10.1016/j.cub.2020.06.058. PubMed PMID: 32679104; PubMed Central PMCID: PMCPMC7484210.

16. Sun C, Wyngaard G, Walton DB, Wichman HA, Mueller RL. Billions of basepairs of recently expanded, repetitive sequences are eliminated from the somatic genome during copepod development. BMC Genomics. 2014;15:186. Epub 20140311. doi: 10.1186/1471-2164-15-186. PubMed PMID: 24618421; PubMed Central PMCID: PMCPMC4029161.

17. Zhao C, Escalante LN, Chen H, Benatti TR, Qu J, Chellapilla S, et al. A massive expansion of effector genes underlies gall-formation in the wheat pest Mayetiola destructor. Curr Biol. 2015;25(5):613-20. Epub 20150205. doi: 10.1016/j.cub.2014.12.057. PubMed PMID: 25660540.

18. Biederman MK, Nelson MM, Asalone KC, Pedersen AL, Saldanha CJ, Bracht JR. Discovery of the first germline-restricted gene by subtractive transcriptomic analysis in the zebra finch, *Taeniopygia guttata*. Curr Biol. 2018;28(10):1620–7 e5. Epub 20180503. doi: 10.1016/j.cub.2018.03.067. PubMed PMID: 29731307; PubMed Central PMCID: PMCPMC5977399.

19. Kinsella CM, Ruiz-Ruano FJ, Dion-Cote AM, Charles AJ, Gossmann TI, Cabrero J, et al. Programmed DNA elimination of germline development genes in songbirds. Nat Commun. 2019;10(1):5468. Epub 20191129. doi: 10.1038/s41467-019-13427-4. PubMed PMID: 31784533; PubMed Central PMCID: PMCPMC6884545.

20. Smith JJ, Timoshevskaya N, Ye C, Holt C, Keinath MC, Parker HJ, et al. The sea lamprey germline genome provides insights into programmed genome rearrangement and vertebrate evolution. Nat Genet. 2018;50(2):270–7. Epub 20180122. doi: 10.1038/s41588-017-0036-1. PubMed PMID: 29358652; PubMed Central PMCID: PMCPMC5805609.

21. Gonzalez de la Rosa PM, Thomson M, Trivedi U, Tracey A, Tandonnet S, Blaxter M. A telomere-to-telomere assembly of *Oscheius tipulae* and the evolution of rhabditid nematode chromosomes. G3 (Bethesda). 2021;11(1). Epub 2021/02/10. doi: 10.1093/g3journal/jkaa020. PubMed PMID: 33561231; PubMed Central PMCID: PMCPMC8022731.

22. Hodson CN, Jaron KS, Gerbi S, Ross L. Gene-rich germline-restricted chromosomes in black-winged fungus gnats evolved through hybridization. PLoS Biol. 2022;20(2):e3001559. Epub 20220225. doi: 10.1371/journal.pbio.3001559. PubMed PMID: 35213540; PubMed Central PMCID: PMCPMC8906591.

23. Wang J. Genomics of the parasitic nematode *Ascaris* and its relatives. Genes (Basel). 2021;12(4). Epub 20210328. doi: 10.3390/genes12040493. PubMed PMID: 33800545; PubMed Central PMCID: PMCPMC8065839.

24. Zagoskin MV, Wang J. Programmed DNA elimination: silencing genes and repetitive sequences in somatic cells. Biochem Soc Trans. 2021;49(5):1891–903. Epub 2021/10/20. doi: 10.1042/BST20190951. PubMed PMID: 34665225; PubMed Central PMCID: PMCPMC9200590.

25. Nagao K, Kubota S, Goto Y. Internal deletion of highly repetitive DNA families from the non-eliminated chromosome in a Japanese hagfish, *Eptatretus burgeri*: First finding in this species. Chromosome Sci. 2021;24(3-4):67–70. doi: 10.11352/scr.24.67.

26. Green MR, Sambrook J. Molecular Cloning: A Laboratory Manual (Fourth Edition): Cold Spring Harbor Laboratory Press; 2012.

27. Pascual-Anaya J, Sato I, Sugahara F, Higuchi S, Paps J, Ren Y, et al. Hagfish and lamprey Hox genes reveal conservation of temporal colinearity in vertebrates. Nat Ecol Evol. 2018;2(5):859–66. Epub 20180402. doi: 10.1038/s41559-018-0526-2. PubMed PMID: 29610468.

28. Staiber W, Wech I, Preiss A. Isolation and chromosomal localization of a germ line-specific highly repetitive DNA family in *Acricotopus lucidus* (Diptera, Chironomidae). Chromosoma. 1997;106(5):267–75. Epub 1997/09/23. doi: 10.1007/s004120050247. PubMed PMID: 9297504.

29. Degtyarev S, Boykova T, Grishanin A, Belyakin S, Rubtsov N, Karamysheva T, et al. The molecular structure of the DNA fragments eliminated during chromatin diminution in *Cyclops kolensis*. Genome Res. 2004;14(11):2287–94. Epub 2004/11/03. doi: 10.1101/gr.2794604. PubMed PMID: 15520291; PubMed Central PMCID: PMCPMC525688.

30. Itoh Y, Kampf K, Pigozzi MI, Arnold AP. Molecular cloning and characterization of the germline-restricted chromosome sequence in the zebra finch. Chromosoma. 2009;118(4):527–36. Epub 20090519. doi: 10.1007/s00412-009-0216-6. PubMed PMID: 19452161; PubMed Central PMCID: PMCPMC2701497.

31. Smith JJ, Antonacci F, Eichler EE, Amemiya CT. Programmed loss of millions of base pairs from a vertebrate genome. Proc Natl Acad Sci U S A. 2009;106(27):11212–7. Epub 20090626. doi: 10.1073/pnas.0902358106. PubMed PMID: 19561299; PubMed Central PMCID: PMCPMC2708698.

32. Bauer H, Beermann W. Der Chromosomencyclus der Orthocladiinen (Nematocera, Diptera). Zeitschrift für Naturforschung B. 1952;7(9-10):557–63. doi: 10.1515/znb-1952-9-1013.

33. Pigozzi MI, Solari AJ. Germ cell restriction and regular transmission of an accessory chromosome that mimics a sex body in the zebra finch, *Taeniopygia guttata*. Chromosome Res. 1998;6(2):105–13. Epub 1998/05/16. doi: 10.1023/a:1009234912307. PubMed PMID: 9543013.

34. Staiber W. Molecular evolution of homologous gene sequences in germline-limited and somatic chromosomes of *Acricotopus*. Genome. 2004;47(4):732–41. Epub 2004/07/31. doi: 10.1139/g04-026. PubMed PMID: 15284878.

35. Torgasheva AA, Malinovskaya LP, Zadesenets KS, Karamysheva TV, Kizilova EA, Akberdina EA, et al. Germline-restricted chromosome (GRC) is widespread among songbirds. Proc Natl Acad Sci U S A. 2019;116(24):11845–50. Epub 20190429. doi: 10.1073/pnas.1817373116. PubMed PMID: 31036668; PubMed Central PMCID: PMCPMC6575587.

36. Metz CW. Chromosome Behavior, Inheritance and Sex Determination in *Sciara*. The American Naturalist. 1938;72(743):485–520. doi: 10.1086/280803.

37. Camacho JP, Sharbel TF, Beukeboom LW. B-chromosome evolution. Philos Trans R Soc Lond B Biol Sci. 2000;355(1394):163-78. Epub 2000/03/21. doi: 10.1098/rstb.2000.0556. PubMed PMID: 10724453; PubMed Central PMCID: PMCPMC1692730.

38. Mestriner CA, Galetti PM, Jr., Valentini SR, Ruiz IR, Abel LD, Moreira-Filho O, et al. Structural and functional evidence that a B chromosome in the characid fish *Astyanax scabripinnis* is an isochromosome. Heredity (Edinb). 2000;85 (Pt 1)(1):1-9. Epub 2000/09/06. doi: 10.1046/j.1365-2540.2000.00702.x. PubMed PMID: 10971685.

39. Silva DMZA, Pansonato-Alves JC, Utsunomia R, Araya-Jaime C, Ruiz-Ruano FJ, Daniel SN, et al. Delimiting the origin of a B chromosome by FISH mapping, chromosome painting and DNA sequence analysis in *Astyanax paranae* (Teleostei, Characiformes). PLoS One. 2014;9(4):e94896. Epub 20140415. doi: 10.1371/journal.pone.0094896. PubMed PMID: 24736529; PubMed Central PMCID: PMCPMC3988084.

40. Silva DMZA, Daniel SN, Camacho JP, Utsunomia R, Ruiz-Ruano FJ, Penitente M, et al. Origin of B chromosomes in the genus *Astyanax* (Characiformes, Characidae) and the limits of chromosome painting. Mol Genet Genomics. 2016;291(3):1407–18. Epub 20160316. doi: 10.1007/s00438-016-1195-y. PubMed PMID: 26984341.

41. Silva DMZA, Utsunomia R, Ruiz-Ruano FJ, Daniel SN, Porto-Foresti F, Hashimoto DT, et al. High-throughput analysis unveils a highly shared satellite DNA library among three species of fish genus *Astyanax*. Sci Rep. 2017;7(1):12726. Epub 20171010. doi: 10.1038/s41598-017-12939-7. PubMed PMID: 29018237; PubMed Central PMCID: PMCPMC5635008.

42. Chen ZQ, Lin CC, Hodgetts RB. Cloning and characterization of a tandemly repeated DNA sequence in the crane family (Gruidae). Genome. 1989;32(4):646–54. doi: 10.1139/g89-493. PubMed PMID: 2806904.

43. Yamada K, Nishida-Umehara C, Matsuda Y. Characterization and chromosomal distribution of novel satellite DNA sequences of the lesser rhea (*Pterocnemia pennata*) and the greater rhea (*Rhea americana*). Chromosome Res. 2002;10(6):513–23. doi: 10.1023/a:1020996431588. PubMed PMID: 12489832.

44. Shang WH, Hori T, Toyoda A, Kato J, Popendorf K, Sakakibara Y, et al. Chickens possess centromeres with both extended tandem repeats and short non-tandem-repetitive sequences. Genome Res. 2010;20(9):1219–28. Epub 20100609. doi: 10.1101/gr.106245.110. PubMed PMID: 20534883; PubMed Central PMCID: PMCPMC2928500.

45. Ishishita S, Tsuruta Y, Uno Y, Nakamura A, Nishida C, Griffin DK, et al. Chromosome size-correlated and chromosome size-uncorrelated homogenization of centromeric repetitive sequences in New World quails. Chromosome Res. 2014;22(1):15–34. Epub 2014/02/18. doi: 10.1007/s10577-014-9402-3. PubMed PMID: 24532185.

46. Habermann FA, Cremer M, Walter J, Kreth G, von Hase J, Bauer K, et al. Arrangements of macro- and microchromosomes in chicken cells. Chromosome Res. 2001;9(7):569–84. doi: 10.1023/a:1012447318535. PubMed PMID: 11721954.

47. Tanabe H, Habermann FA, Solovei I, Cremer M, Cremer T. Non-random radial arrangements of interphase chromosome territories: evolutionary considerations and functional implications. Mutat Res. 2002;504(1-2):37–45. doi: 10.1016/s0027-5107(02)00077-5. PubMed PMID: 12106644.

48. Alkhimova OG, Mazurok NA, Potapova TA, Zakian SM, Heslop-Harrison JS, Vershinin AV. Diverse patterns of the tandem repeats organization in rye chromosomes. Chromosoma. 2004;113(1):42–52. Epub 20040715. doi: 10.1007/s00412-004-0294-4. PubMed PMID: 15257465.

49. Kuhn GC, Sene FM, Moreira-Filho O, Schwarzacher T, Heslop-Harrison JS. Sequence analysis, chromosomal distribution and long-range organization show that rapid turnover of new and old pBuM satellite DNA repeats leads to different patterns of variation in seven species of the *Drosophila buzzatii* cluster. Chromosome Res. 2008;16(2):307–24. Epub 20080211. doi:10.1007/s10577-007-1195-1. PubMed PMID: 18266060.

50. Kuhn GC, Teo CH, Schwarzacher T, Heslop-Harrison JS. Evolutionary dynamics and sites of illegitimate recombination revealed in the interspersion and sequence junctions of two nonhomologous satellite DNAs in cactophilic *Drosophila* species. Heredity (Edinb). 2009;102(5):453–64. Epub 20090304. doi: 10.1038/hdy.2009.9. PubMed PMID: 19259119.

51. de Barros AV, Sczepanski TS, Cabrero J, Camacho JPM, Vicari MR, Artoni RF. Fiber FISH reveals different patterns of high-resolution physical mapping for repetitive DNA in fish. Aquaculture. 2011;322–323:47-50. doi: 10.1016/j.aquaculture.2011.10.002.

52. Paco A, Adega F, Mestrovic N, Plohl M, Chaves R. The puzzling character of repetitive DNA in *Phodopus* genomes (Cricetidae, Rodentia). Chromosome Res. 2015;23(3):427–40. doi: 10.1007/s10577-015-9481-9. PubMed PMID: 26281779.

53. Milani D, Ramos E, Loreto V, Marti DA, Cardoso AL, de Moraes KCM, et al. The satellite DNA AflaSAT-1 in the A and B chromosomes of the grasshopper *Abracris flavolineata*. BMC Genet. 2017;18(1):81. Epub 20170829. doi: 10.1186/s12863-017-0548-9. PubMed PMID: 28851268; PubMed Central PMCID: PMCPMC5575873.

54. Viana PF, Ezaz T, Marajo L, Ferreira M, Zuanon J, Cioffi MB, et al. Genomic organization of repetitive DNAs and differentiation of an XX/XY sex chromosome system in the Amazonian puffer fish, *Colomesus asellus* (Tetraodontiformes). Cytogenet Genome Res. 2017;153(2):96–104. Epub 20171130. doi: 10.1159/000484423. PubMed PMID: 29186711.

55. Souza J, Guimaraes E, Pinheiro-Figliuolo V, Cioffi MB, Bertollo LAC, Feldberg E. Chromosomal Analysis of Ctenolucius hujeta Valenciennes, 1850 (Characiformes): A new piece in the chromosomal evolution of the Ctenoluciidae. Cytogenet Genome Res. 2021;161(3-4):195-202. Epub 20210614. doi: 10.1159/000515456. PubMed PMID: 34126615.

